# Increased green autofluorescence is a marker for non-invasive prediction of H_2_O_2_-induced cell death and decreases in the intracellular ATP of HaCaT cells

**DOI:** 10.1101/298075

**Authors:** Jie Xu, Weihai Ying

**Affiliations:** Med-X Research Institute and School of Biomedical Engineering, Shanghai Jiao Tong University, Shanghai 200030, P.R. China; Collaborative Innovation Center for Genetics and Development, Shanghai 200043, P.R. China

**Keywords:** Oxidative stress, autofluorecence, apoptosis, necrosis, ATP, keratinocytes

## Abstract

Since oxidative stress plays important pathological roles in numerous diseases, it is of both critical theoretical and clinical significance to search for the approaches for predicting oxidative damage. Cellular models have great value for studying oxidative damage, which would be significantly promoted if non-invasive approaches for predicting oxidative damage can be established without the need of exogenous probes. In our current study, we tested our hypothesis that changes of the autofluorescence (AF) of cells may be used for predicting oxidative cellular damage. Our study found that H_2_O_2_ dose-dependently increased the green AF of HaCaT keratinocyte cell line at non-nuclear regions assessed at 1 hr or 3 hrs after the H_2_O_2_ exposures, while H_2_O_2_ did not affect the green AF of other cell types tested in our study, including PC 12 cells and BV2 microglia. We further found that the increases in the AF of HaCaT cells are highly correlated with the H_2_O_2_-induced increases in early-stage apoptosis, late-stage apoptosis and necrosis assessed at 18 hrs after the H_2_O_2_ exposures, which are also negatively correlated with the intracellular ATP levels of the H_2_O_2_-treated cells assessed at 18 hrs after the H_2_O_2_ exposures. Collectively, our study has suggested that increased AF may become the first endogenous marker for non-invasive prediction of oxidative damage selectively for such cell types as HaCaT cells. Compared with traditional approaches, our method may have significant value for studying oxidative damage of keratinocytes with significantly higher efficiency and lower cost.

## Introduction

Oxidative stress plays critical roles in aging and multiple diseases including cerebra ischemia (8), myocardial ischemia (15), Alzheimer’s disease (4), head trauma (17) and diabetes (7). Prevention of oxidative damage has been an important strategy for both decreasing the pathological changes of multiple diseases and slowing the aging process. It is both critical theoretical and clinical significance to search for the approaches for monitoring oxidative stress and predicting oxidative damage in cells and tissues.

Cellular models are highly valuable for studying both the mechanisms underlying oxidative damage and the antioxidant capacity of molecules. Currently, there are two major approaches for monitoring intracellular oxidative stress: The non-invasive approaches require applications of such fluorescent probes as dichlorofluorescein (DCF) (18) and dihydroethidium (DHE) (14); while the other approaches used for either determining antioxidant capacity of cells (3) or assessing oxidative damage (19) require cell lysis. There has been no non-invasive approaches that can predict oxidative cellular damage.

Human AF (AF) of skin or blood has been used as biomarkers for non-invasive diagnosis of diabetes (12) and cancer (16). Our recent study has also suggested that UV-induced epidermal green AF can be used as a novel biomarker for predicting UV-induced skin damage (9). Our study has also suggested that UV-induced keratin 1 proteolysis mediates the UV-induced increases in the epidermal AF (9). However, it is unknown if AF may be used in cellular models for non-invasive predication of oxidative stress-induced apoptosis, necrosis and other cellular alterations.

The major goal of our current study is to test our hypothesis that oxidative stress-induced AF changes of cells may be used for non-invasive prediction of oxidative cellular damage. Our study has found that H_2_O_2_ can dose-dependently induce increases in the green AF of HaCaT cells at 1 hr and 3 hrs after the H_2_O_2_ exposures, which are highly correlated with the H_2_O_2_-induced early-stage apoptosis, late-stage apoptosis and necrosis assessed at 18 hrs after the H_2_O_2_ exposures. The AF changes are also highly negatively correlated with the intracellular ATP levels of the cells. In contrast, H_2_O_2_ did not change the green AF of other cell types tested in our study, including PC 12 cells and BV2 microglia. Therefore, the increased green AF may become the first endogenous marker for non-invasive prediction of oxidative cellular damage of such cell types as HaCaT cells.

## Methods and Materials

### Cell cultures

HaCaT cells, BV2 microglia or PC12 cells were plated into 24-well or 12-well cell culture plates at the initial density of 1 × 10^6^ cells/mL in Dulbecco’s Modified Eagle’s Medium containing 4500 mg/L D-glucose, 584 mg/L L-glutamine (Thermo Scientific, Waltham, MA, USA), and 1% penicillin and streptomycin (Invitrogen, Carlsbad, CA, USA), supplemented with 10% fetal bovine serum (PAA, Germany). The cells were maintained in a 5% CO_2_ incubator at 37 °C.

### 2. Autofluorescence imaging of HaCaT cells

After treatment of cells with H_2_O_2_, the cells were washed with PBS and re-suspended in primary DMEM medium containing 10% FBS. One hr or 3 hours after the H_2_O_2_ treatment, the AF signals of the cells were observed under a Leica confocal fluorescence microscope at the excitation wavelength of 488 nm and the emission wavelength between 500 to 530 nm.

### 3. Hoechst staining

The nuclei of HaCaT cells were stained with Hoechst 33342 Staining Solution for Live Cells (Beyotime, Jiangsu Province, China) for 10 min. After two washes with PBS, the cells were photographed under a Leica confocal fluorescence microscope. The excitation wavelength was 350 nm and the emission wavelength was at the range between 450 to 470 nm.

### 4. Intracellular lactate dehydrogenase (LDH) assay

As described previously (11), cell survival was quantified by measuring the intracellular LDH activity of the cells. In brief, cells were lysed for 15 min in lysing buffer containing 0.04% Triton X-100, 2 mM HEPES and 0.01% bovine serum albumin (pH 7.5). Then 50 μL cell lysates were mixed with 150 μL 500 mM potassium phosphate buffer (pH 7.5) containing 0.34 mM NADH and 2.5 mM sodium pyruvate. The A340nm changes were monitored over 90 sec. Percentage of cell survival was calculated by normalizing the LDH values of samples to LDH activity measured in the lysates of control (wash only) culture wells.

### 5. FACS-based Annexin V/7-AAD assay

As described previously (10), FACS-based Annexin V/7-AAD assay was performed to determine the levels of early-stage apoptosis, late-stage apoptosis and necrosis by using ApoScreen Annexin V kit (SouthernBiotech, Birmingham, AL, USA) according to the manufacturer’s protocol. In brief, cells were digested with 0.25% trypsin-EDTA, washed by cold PBS one time and resuspended in cold 1X binding buffer (10 mM HEPES, pH 7.4, 140 mM NaCl, 2.5 mM CaCl_2_, 0.1% BSA) at concentrations between 1 × 10^6^ and 1 × 10^7^ cells/mL. Then 5 μL of labeled Annexin V was added into 100 μL of the cell suspension. After incubation on ice for 15 min, 200 μL 1X binding buffer and 5 μL 7-AAD solution were added into the cell suspensions. The number of stained cells was assessed immediately by a flow cytometer (FACSAria II, BD Biosciences).

### 6. ATP assay

As described previously (5), intracellular ATP levels were determined using an ATP Bioluminescence Assay Kit (Roche Applied Science, Mannheim, Germany) following the standard protocol provided by the vendor. Briefly, the cells were lysed with the Cell Lysis Reagent on ice, and 50 μL of the lysates was mixed with 50 μL of the Luciferase Reagent. Then the chemiluminescence of the samples was measured by using a plate reader. The ATP concentrations of the samples were calculated using ATP standard, and normalized to the protein concentrations of the samples, which were determined using the Pierce^®^ BCA Protein Assay Kit (Thermo Fisher Scientific, USA).

### 7. Statistical analyses

All data are presented as mean ± SEM. Data were assessed by one-way ANOVA, followed by Student - Newman - Keuls *post hoc* test. P values less than 0.05 were considered statistically significant

## Results

We determined if oxidative stress may induce changes of the green AF of HaCaT cells that have been widely used for studies on oxidative damage of skin cells (1, 20). We found that treatment of the cells with 0.1, 0.3, or 0.5 mM H_2_O_2_ for 15 minutes led to obvious increases in green AF, assessed at 1 hr or 3 hrs after the H_2_O_2_ exposures (Fig. 1A). Quantifications of the AF indicate that H_2_O_2_ produced dose-dependent increases in the AF (Fig. 1B). By staining the nuclei of HaCaT cells by Hoechst 33342, we found that virtually none of the green AF was observed at the nuclei, suggesting that the AF increases occur mainly at non-nuclear regions of the cells. We also determined the effects of H_2_O_2_ on the green AF of BV2 microglia (Supplemental Fig. 1A and 1B) and PC 12 cells - a neuron-like cell line (Supplemental Fig. 1C and 1D), showing that H_2_O_2_ was incapable of affecting the green AF of these two cell types.

**Figure 1.**
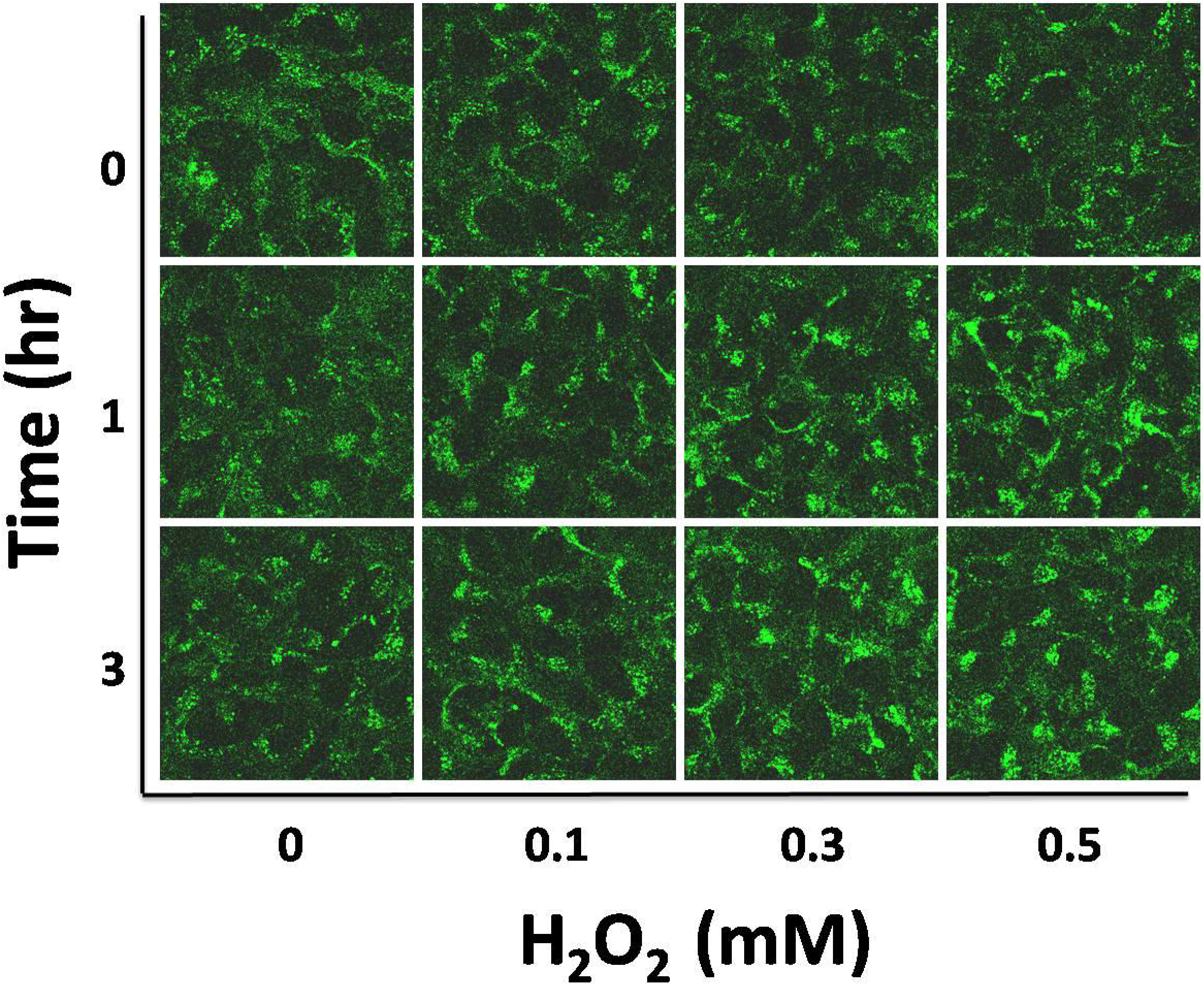
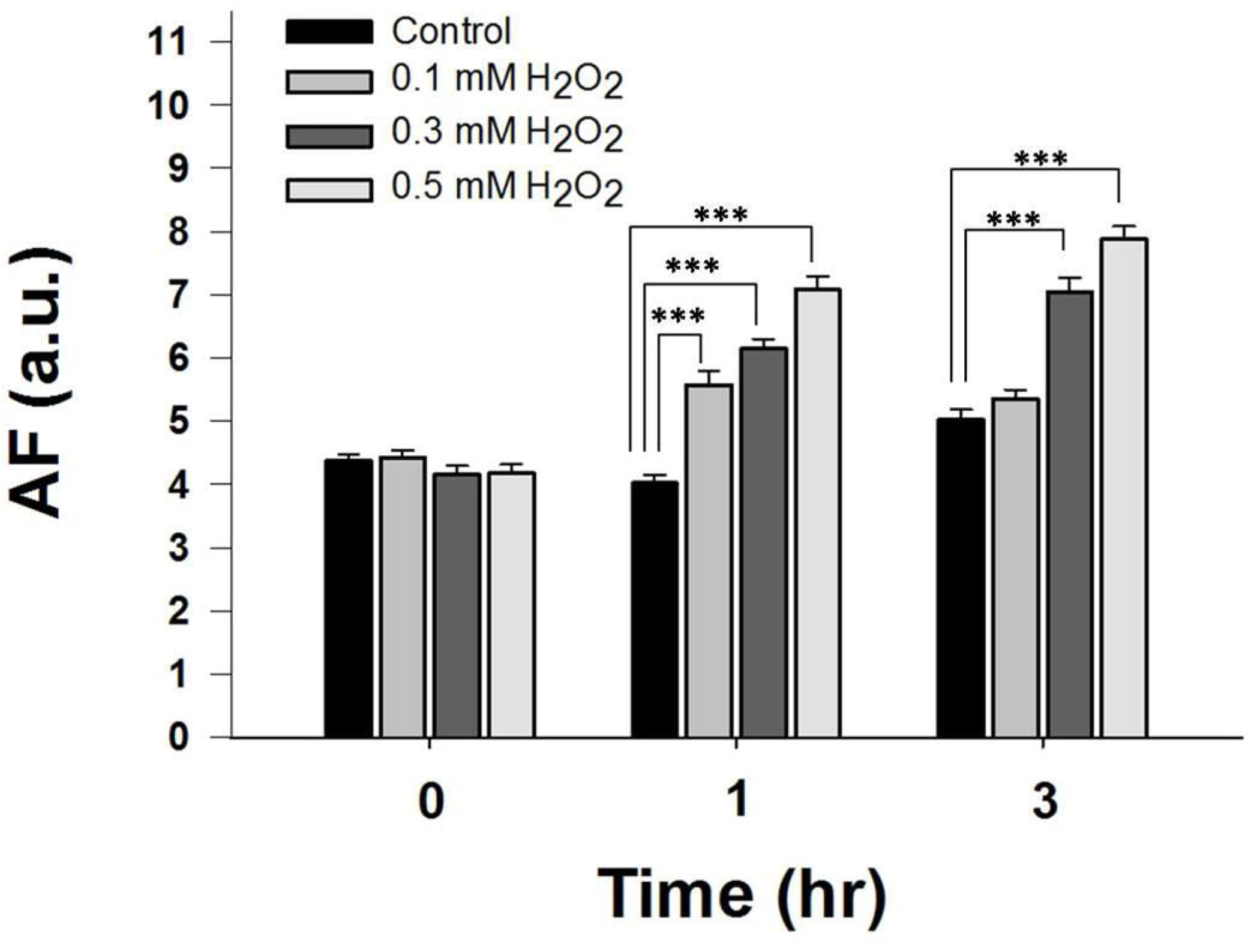
H_2_O_2_ induced increases in the green AF of HaCaT cell. (A) Treatment of HaCaT cells with H_2_O_2_ led to increases in the AF of HaCaT cells. HaCaT cells were treated by 0.1, 0.3, or 0.5 mM H_2_O_2_ for 15 mins. The green AF of the cells was determined under a confocal microscope at 1 hr or 3 hrs after the H_2_O_2_ exposures. The exciting wavelength was 488 nm and the emission wavelength was at the range between 500 and 530 nm. (B) Quantifications of the AF show that H_2_O_2_ dose-dependently induced significant increases in the AF. N = 9. Data were collected from three independent experiments. \**p* < 0.05; \*\*\**p* < 0.001.

We conducted FACS-based assay to determine the effects of H_2_O_2_ on the early-stage apoptosis, late-stage apoptosis and necrosis of HaCaT cells 18 hours after treatment of the cells with 0.1, 0.3, or 0.5 mM H_2_O_2_. H_2_O_2_ led to increases in early-stage apoptosis, late-stage apoptosis and necrosis of HaCaT cells (Fig. 3A). Quantifications of the results indicate that H_2_O_2_ dose-dependently induced significant increases both in early-stage apoptosis and total cell death - the sum of the cells in early-stage apoptosis, late-stage apoptosis and necrosis (Fig. 3B). We further found that the H_2_O_2_-induced AF at 1 hr after the H_2_O_2_ exposures is highly positively correlated with the H_2_O_2_-induced increases in early-stage apoptosis (Fig. 3C), late-stage apoptosis (Fig. 3D), necrosis (Fig. 3E) and total cell death (Fig. 3F), which was assessed at 18 hrs after the H_2_O_2_ exposures. We also found that the H_2_O_2_-induced green AF at 1 hr after the H_2_O_2_ exposures is highly negatively correlated with the cell survival of H_2_O_2_-treated HaCaT cells (Supplemental Figs. 2A and 2B).

**Figure 2.**
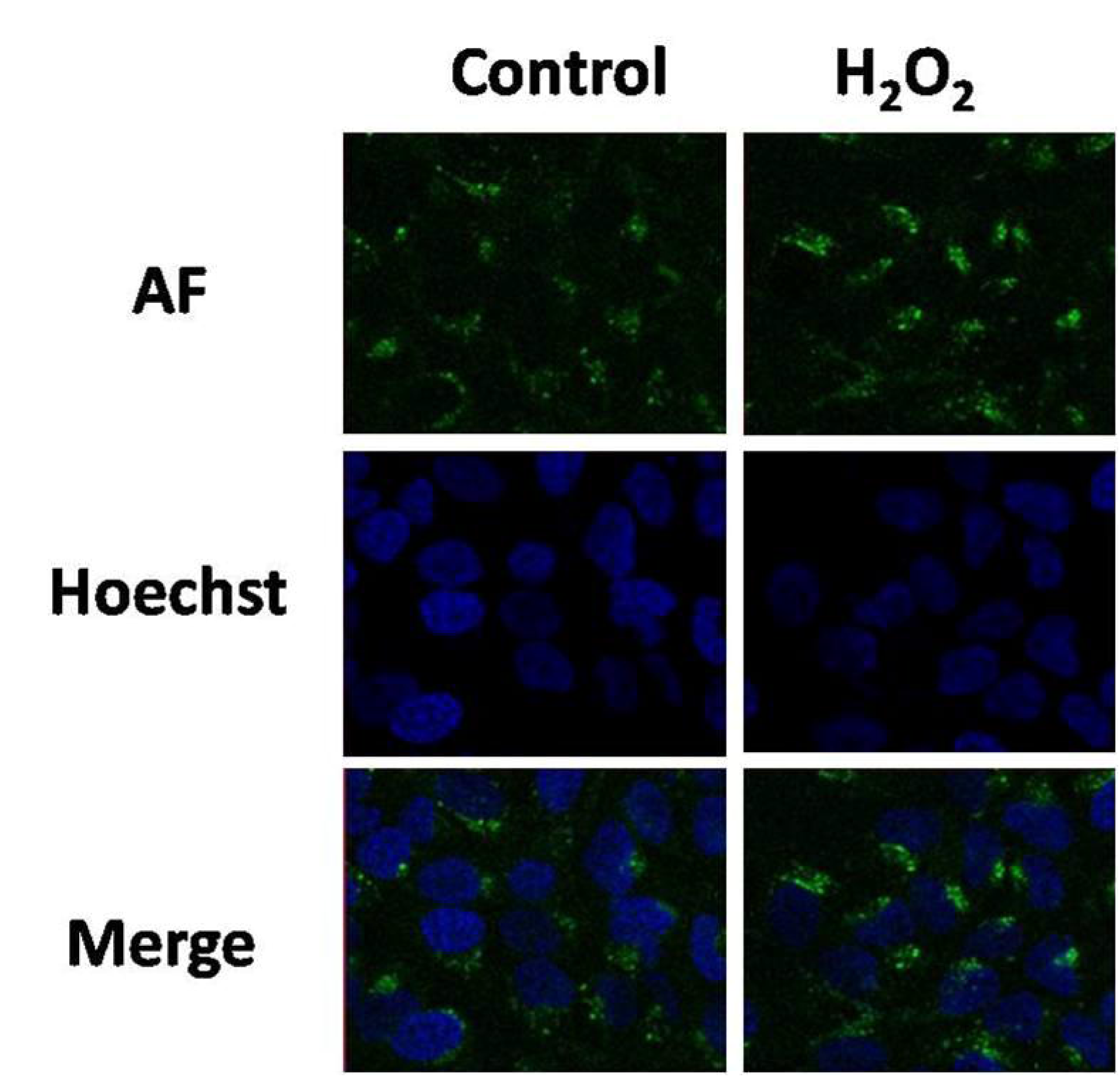
H_2_O_2_ induced increases in the AF at the non-nuclear regions of HaCaT cells. After the nuclei of H_2_O_2_-treated HaCaT cells were stained with Hoechst 33342 for 10 min, the cells were examined under a confocal fluorescence microscope. For determining the fluorescence of Hoechst 33342, the exciting wavelength was 350 nm and the emission wavelength was at the range between 450 - 470 nm. For determining the green AF, the exciting wavelength was 488 nm and the emission wavelength was at the range between 500 and 530 nm.

**Figure 3.**
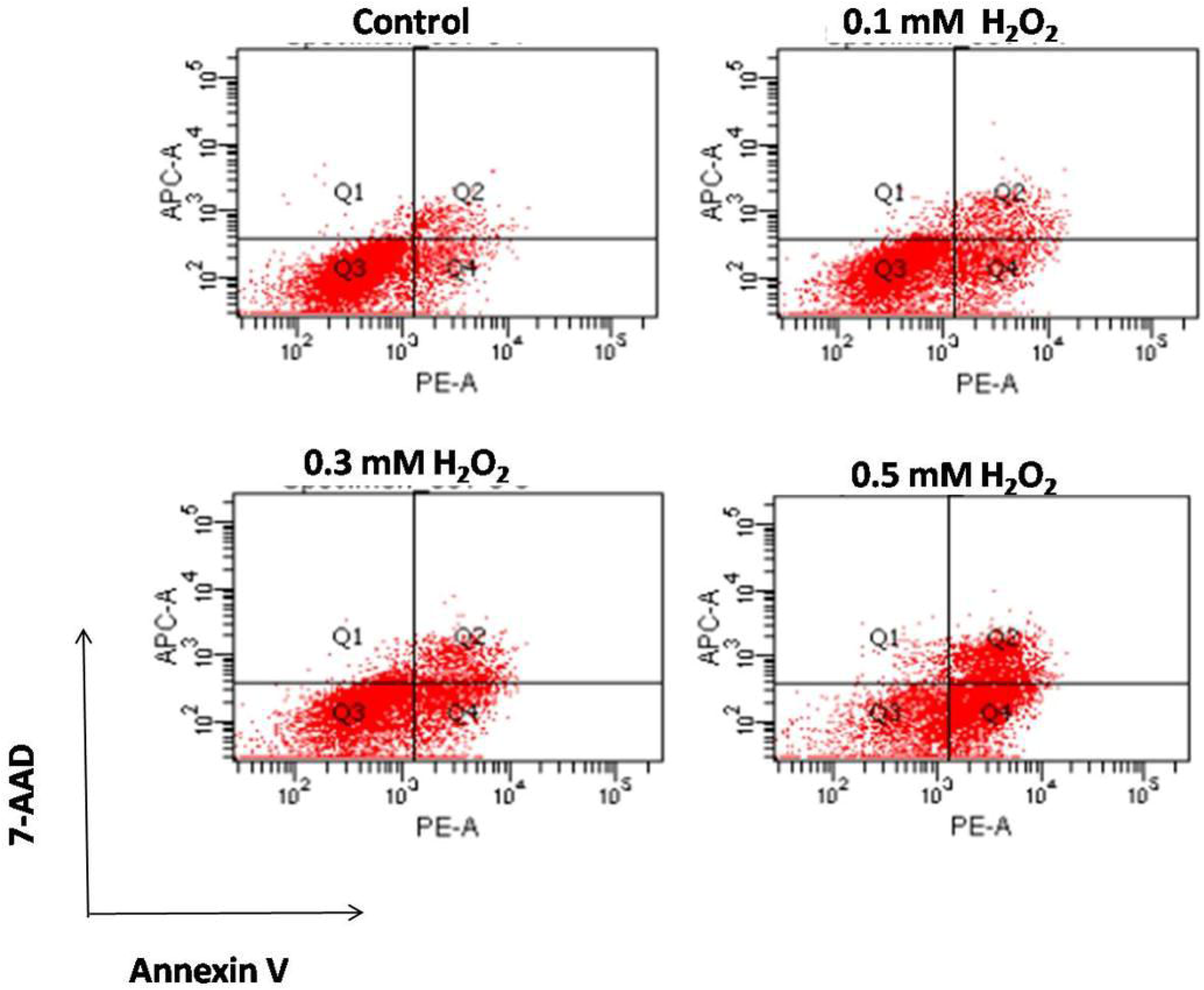
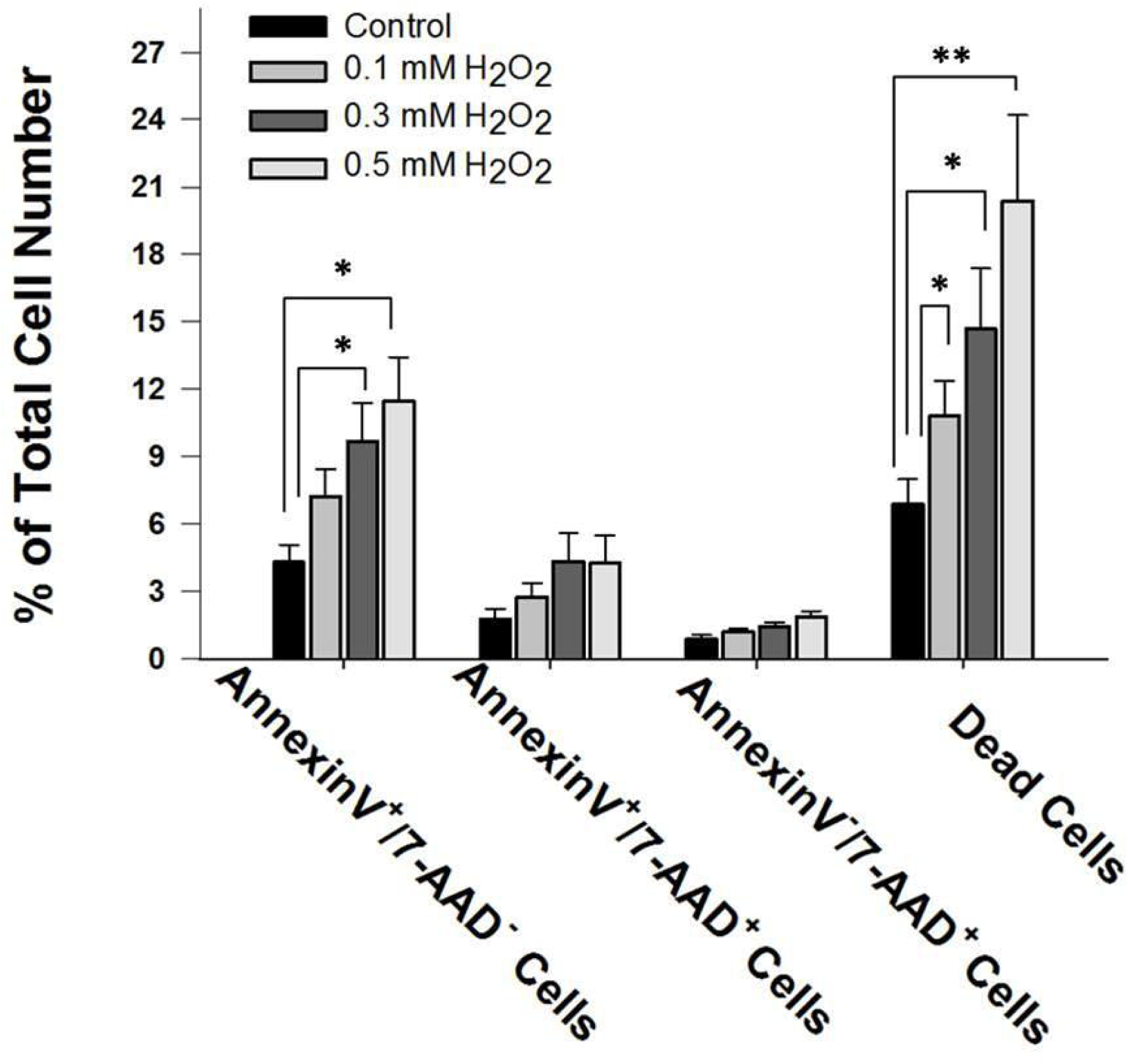
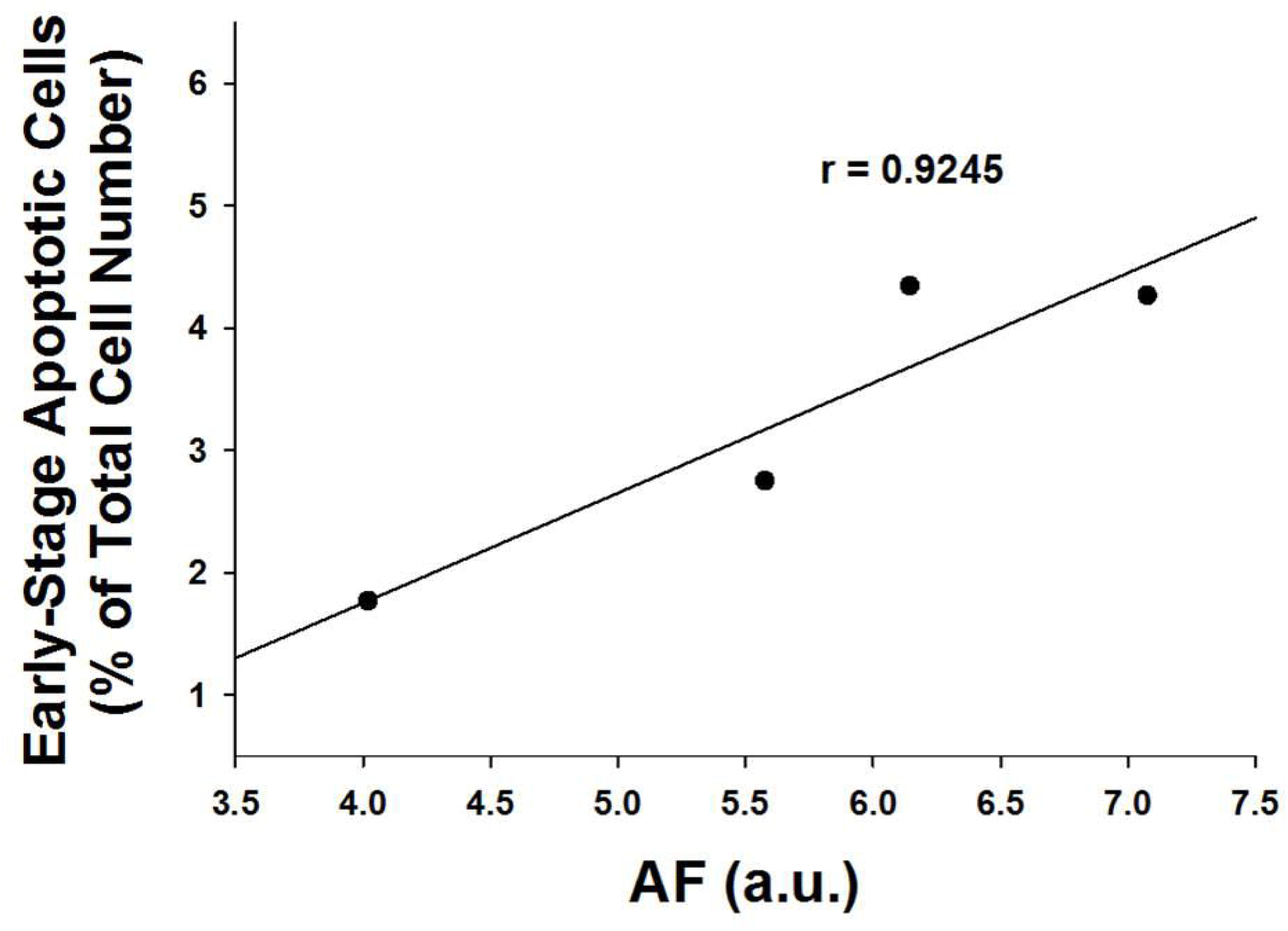
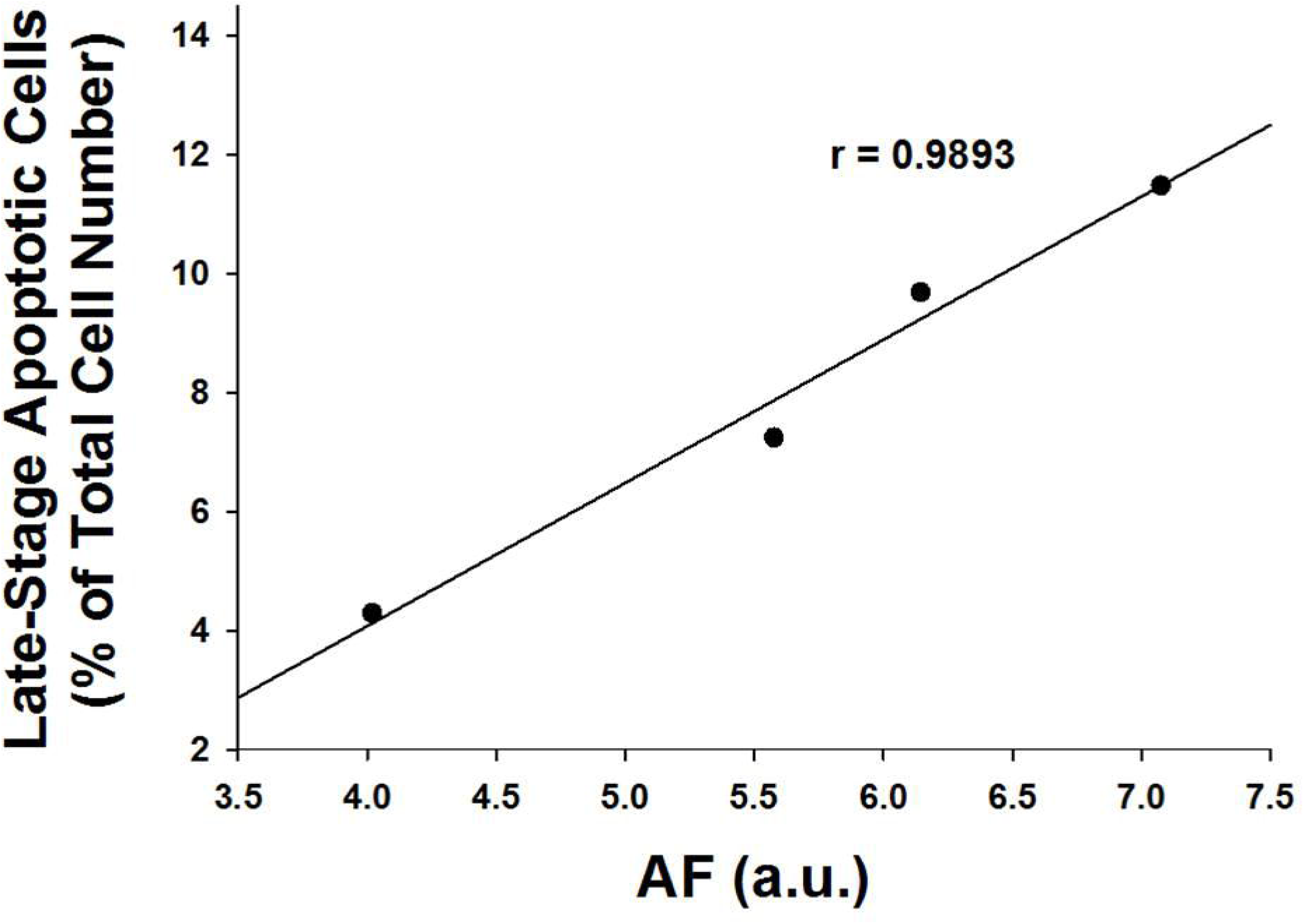
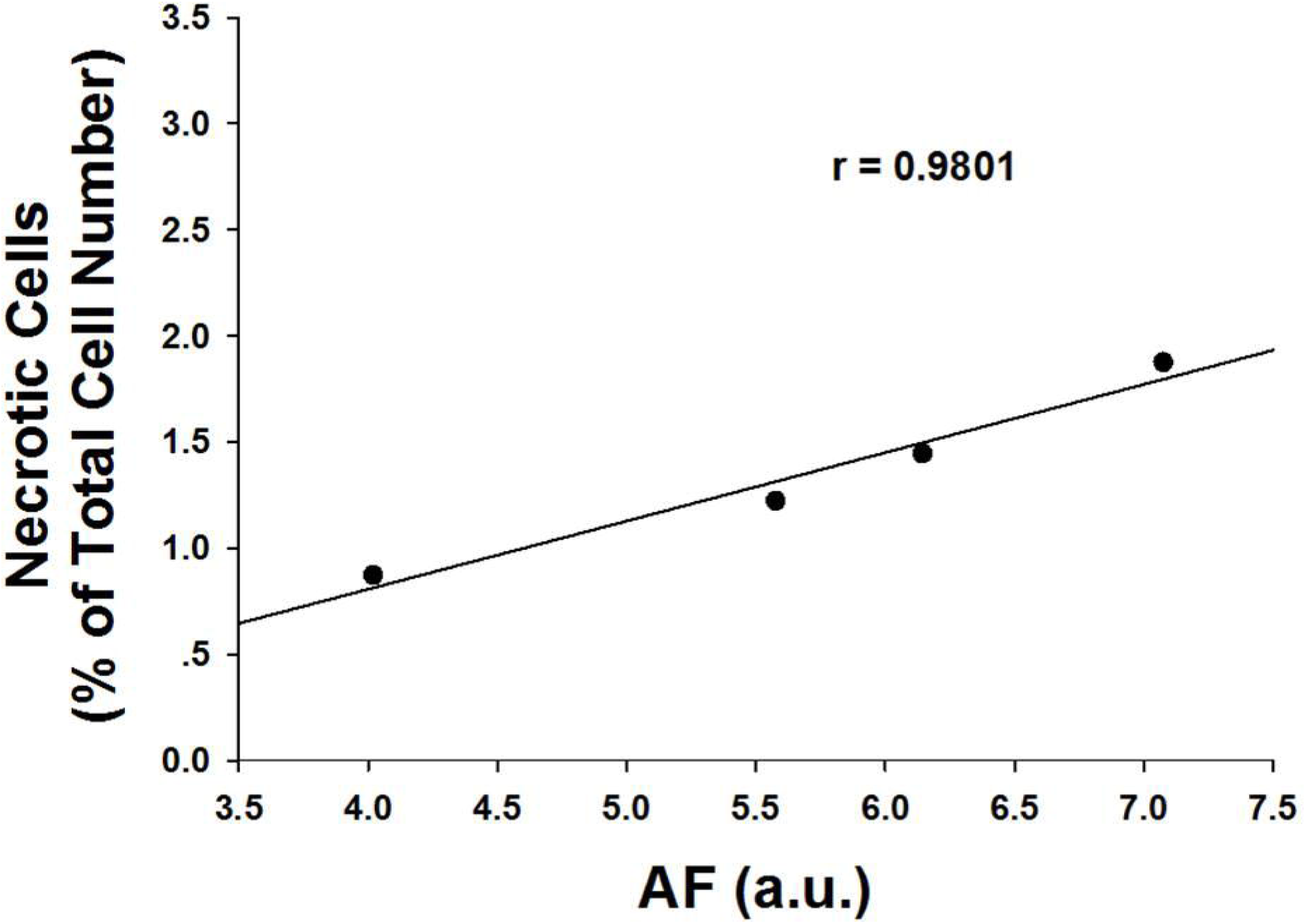
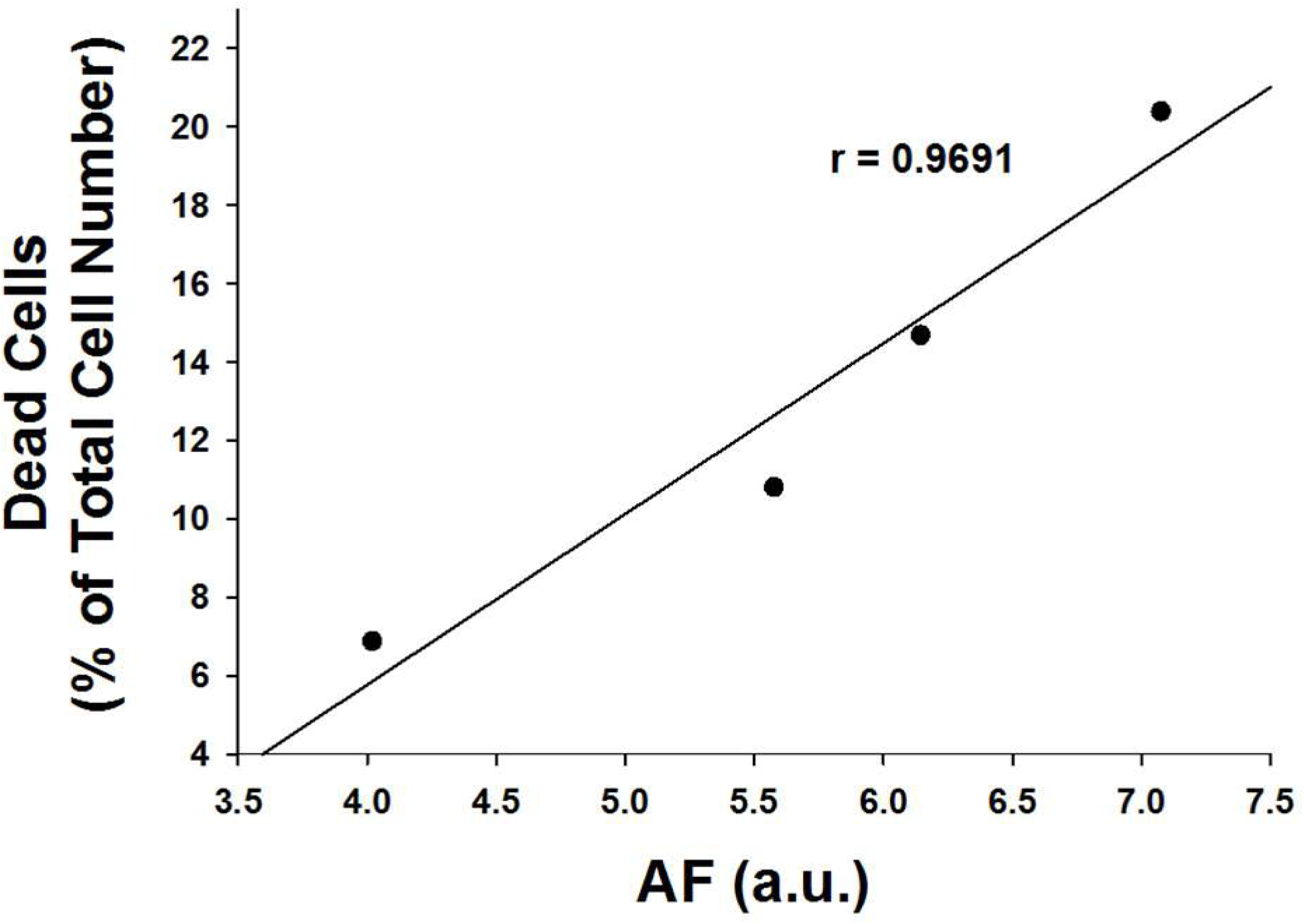
H_2_O_2_-induced green AF at 1 hr after H_2_O_2_ exposures is highly positively correlated with the H_2_O_2_-induced early-stage apoptotic cell death, late-stage apoptotic cell death, necrosis and total cell death at 18 hrs after H_2_O_2_ exposures. (A) FACS-based assay showed that treatment of HaCaT cells with H_2_O_2_ led to obvious increases in cell death of HaCaT cells at 18 hours after H_2_O_2_ treatment. (B) Quantifications of the cell death have indicated that H_2_O_2_ induced significant increases in both early-stage apoptosis and total cell death - the sum of early-stage apoptosis, late-stage apoptosis and necrosis. N = 9. Data were collected from three independent experiments. \**p* < 0.05; \*\*\**p* < 0.001. (C) The H_2_O_2_-induced green AF at 1 hr after H_2_O_2_ exposures is highly positively correlated with the H_2_O_2_-induced early-stage apoptotic cell death at 18 hrs after H_2_O_2_ exposures. (D) The H_2_O_2_-induced green AF at 1 hr after H_2_O_2_ exposures is highly positively correlated with the H_2_O_2_-induced late-stage apoptotic cell death at 18 hrs after H_2_O_2_ exposures. (E) The H_2_O_2_-induced green AF at 1 hr after H_2_O_2_ exposures is highly positively correlated with the H_2_O_2_-induced necrosis at 18 hrs after H_2_O_2_ exposures. (F) The H_2_O_2_-induced green AF at 1 hr after H_2_O_2_ exposures is highly positively correlated with the H_2_O_2_-induced total cell death at 18 hrs after H_2_O_2_ exposures.

We also determined the effects of H_2_O_2_ on the intracellular ATP levels of HaCaT cells 18 hours after H_2_O_2_ treatment. H_2_O_2_ led to obvious decreases in the intracellular ATP levels (Fig. 4A). Quantifications of the results indicate that H_2_O_2_ dose-dependently induced decreases in the intracellular ATP levels of the cells (Fig. 4A). The H_2_O_2_-induced AF at 1 hr after the H_2_O_2_ exposures is highly negatively correlated with the intracellular ATP levels in the H_2_O_2_-treated cells assessed at 18 hrs after the H_2_O_2_ exposures (Fig. 4B).

**Figure 4.**
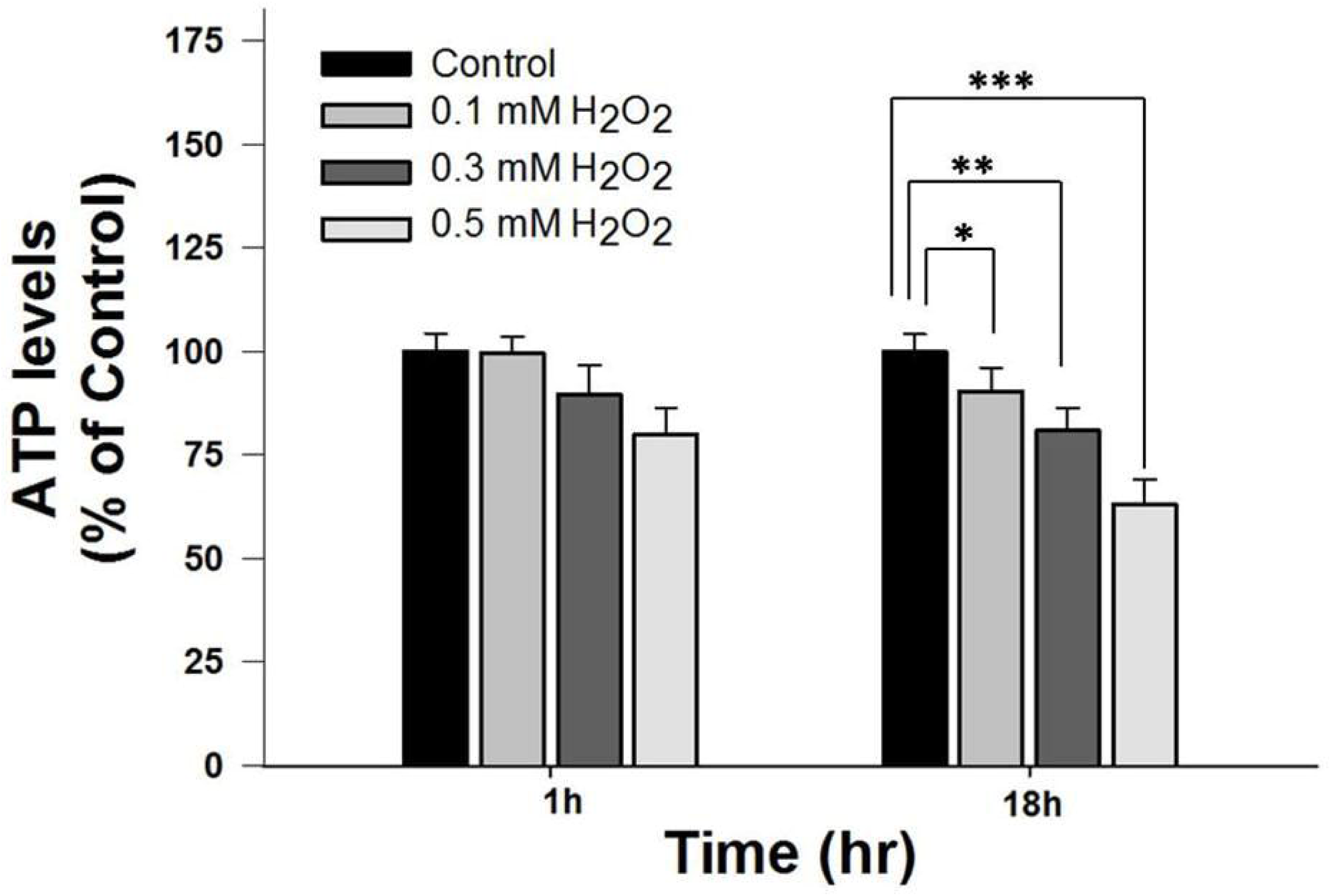
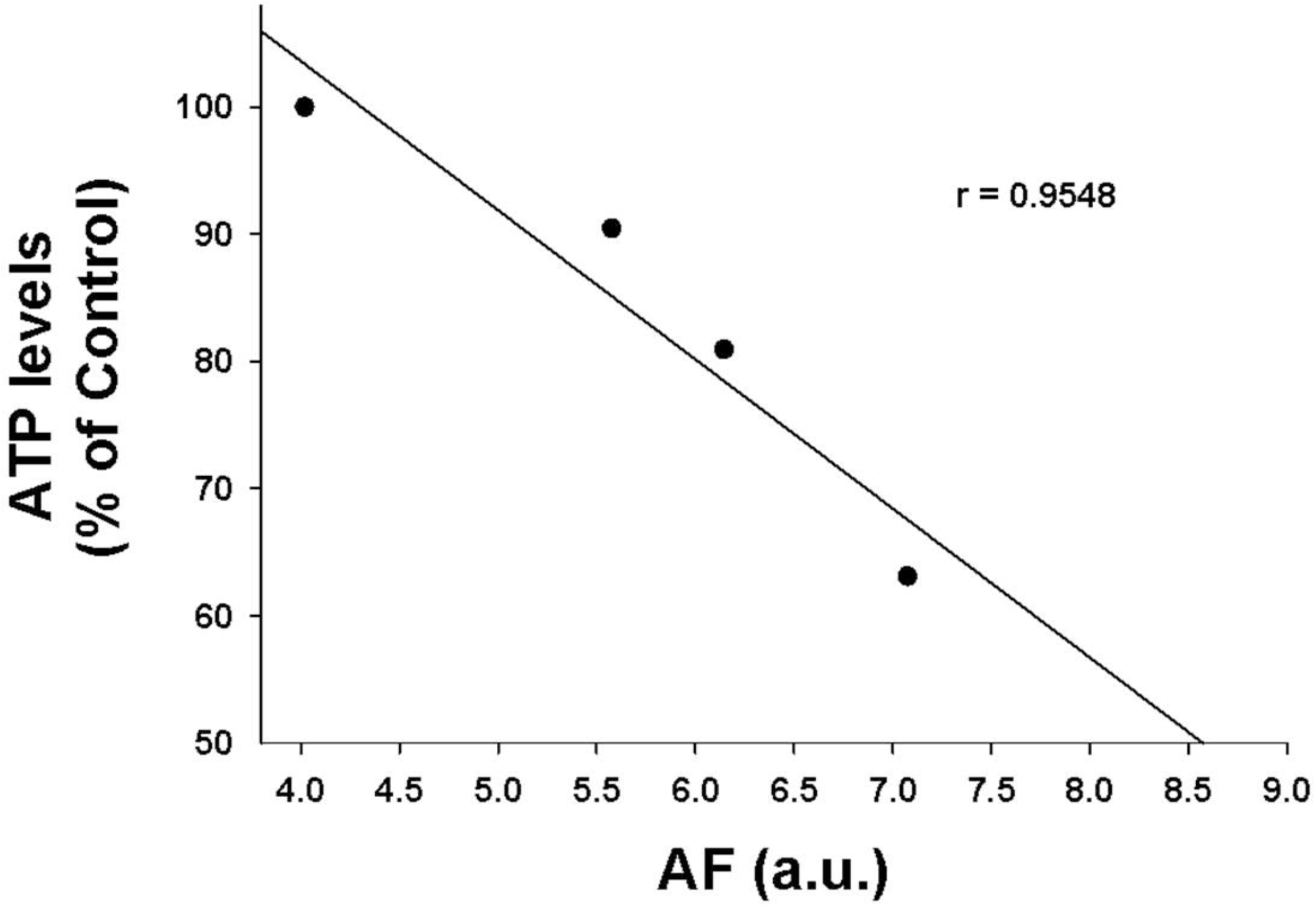
H_2_O_2_-induced AF at 1 hr after H_2_O_2_ exposures is highly negatively correlated with the intracellular ATP levels of H_2_O_2_-treated HaCaT cells at 18 hrs after H_2_O_2_ exposures. (A) FACS-based assay showed that treatment of HaCaT cells with H_2_O_2_ led to obvious decreases in the intracellular ATP levels at 18 hrs after H_2_O_2_ treatment. N = 9. Data were collected from three independent experiments. \**p* < 0.05; \*\*\**p* < 0.001. (B) H_2_O_2_-induced AF at 1 hr after H_2_O_2_ exposures is highly negatively correlated with the intracellular ATP levels of H_2_O_2_-treated HaCaT cells at 18 hrs after H_2_O_2_ exposures.

## Discussion

The major findings of our study include: First, H_2_O_2_ can induce dose-dependent increases in green AF of HaCaT keratinocytes at non-nuclear regions at both 1 and 3 hrs after H_2_O_2_ exposures; second, the H_2_O_2_-induced increases in the green AF of HaCaT cells at early stage of the H_2_O_2_ exposures are highly positively correlated with the H_2_O_2_-induced early-stage apoptosis, late-stage apoptosis, necrosis and total cell death of HaCaT cells at late stage of the H_2_O_2_ exposures; and third, the H_2_O_2_-induced increases in the green AF of HaCaT cells at early stage of H_2_O_2_ exposures are highly negatively correlated with the intracellular ATP levels in the H_2_O_2_-treated HaCaT cells assessed at late stage of the H_2_O_2_ exposures.

Our study has shown that H_2_O_2_ can induce dose-dependent increases in the green AF of HaCaT cells. In contrast to previous reports that oxidative stress can lead to decreased green AF of NAD(P)H of rat hepatocytes (2), our current study shows that oxidative stress can induce increased green AF of HaCaT cells. Our previous study has indicated that UV, a factor that can induce oxidative stress, induces increased green AF of mouse skin by producing cysteine protease-mediated degradation of keratin 1 (9). Because HaCaT keratinocytes express keratin 1 (6,13), we propose that H_2_O_2_ may induce increased green AF of the cells by producing alterations of keratin 1. Future studies are needed to test the validity of this proposal.

Our study has also indicated that the H_2_O_2_-induced increases in the green AF in HaCaT cells at early stage of H_2_O_2_ exposures are highly positively correlated with H_2_O_2_-induced early-stage apoptosis, late-stage apoptosis, necrosis and total cell death. The H_2_O_2_-induced increases in the green AF are also highly negatively correlated with the intracellular ATP levels of HaCaT cells at late stage of H_2_O_2_ exposures. Therefore, our study has suggested that the H_2_O_2_-induced increases in the green AF could become the first endogenous marker for predicting oxidative cell death and loss of intracellular ATP levels of such cell types as HaCaT cells.

Because oxidative stress plays critical roles in numerous diseases and cell death under many situations, our AF-based method for predicting oxidative damage of HaCaT cells may have significant applications for studying oxidative damage of keratinocytes: For studying the effects of certain drugs or other factors on oxidative stress in HaCaT cells, our method can rapidly determine if the factors can change oxidative stress-induced cell death with much higher efficiency and lower cost, compared with traditional approaches. Because HaCaT is a widely used keratinocyte cell line, our study has provided a novel and efficient approach for studying oxidative damage of keratinocyte.

Our study has indicated that hydrogen peroxide can induce the increases in green AF selectively in such cell types as HaCaT cells, but not in the other cell types tested in our study, including PC12 cells and BV2 microglia. The mechanisms underlying these observations require future investigation. Future studies are also needed to determine which types of cells also possess the property that oxidative stress can induce increases in the green AF of the cells.

## Acknowledgments

The authors would like to acknowledge the financial support by a Major Research Grant from the Scientific Committee of Shanghai Municipality #16JC1400502 (to W.Y.) and Chinese National Natural Science Foundation Grant #81271305 (to W. Y.).

---

**Supplemental Figure 1.**
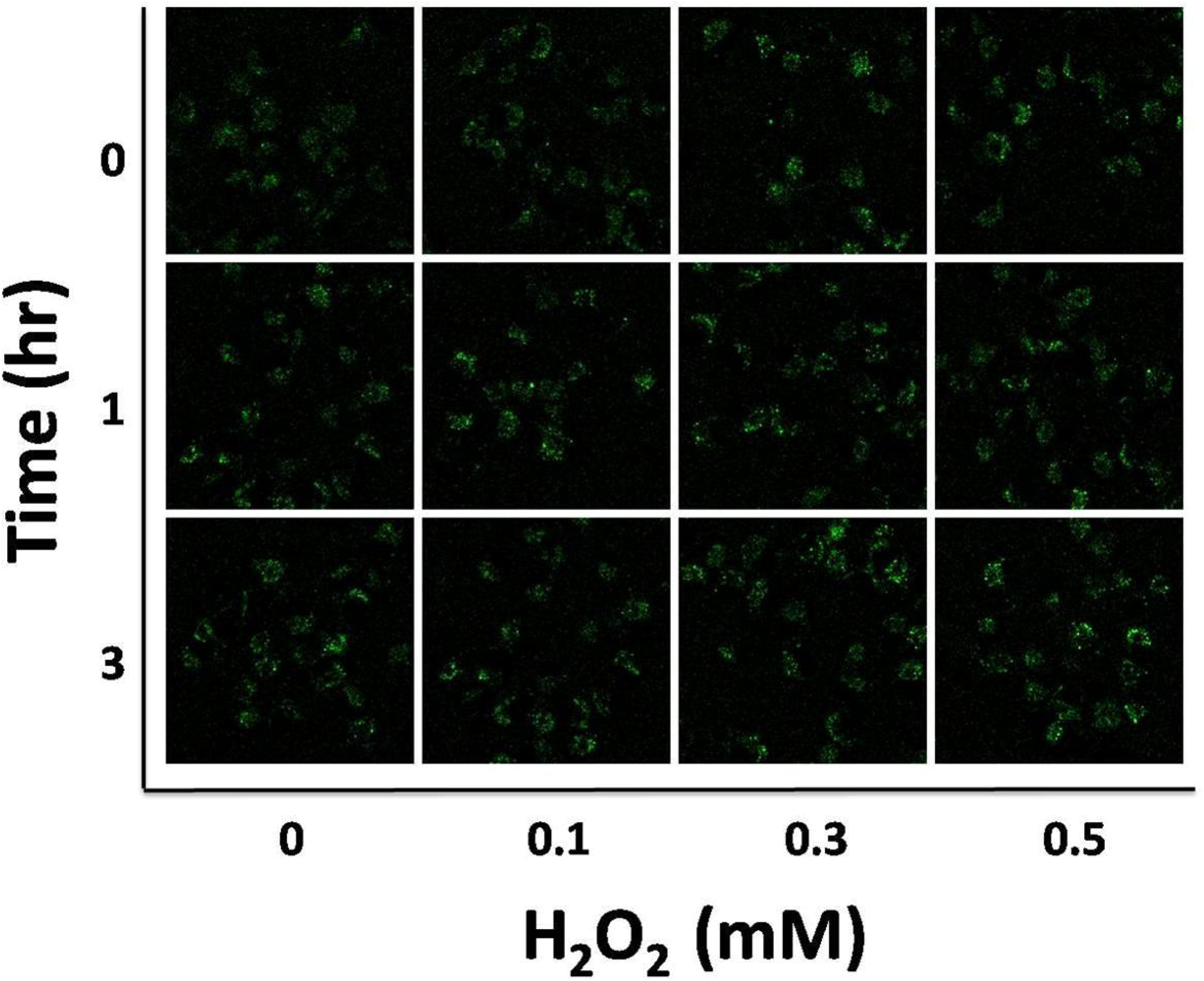
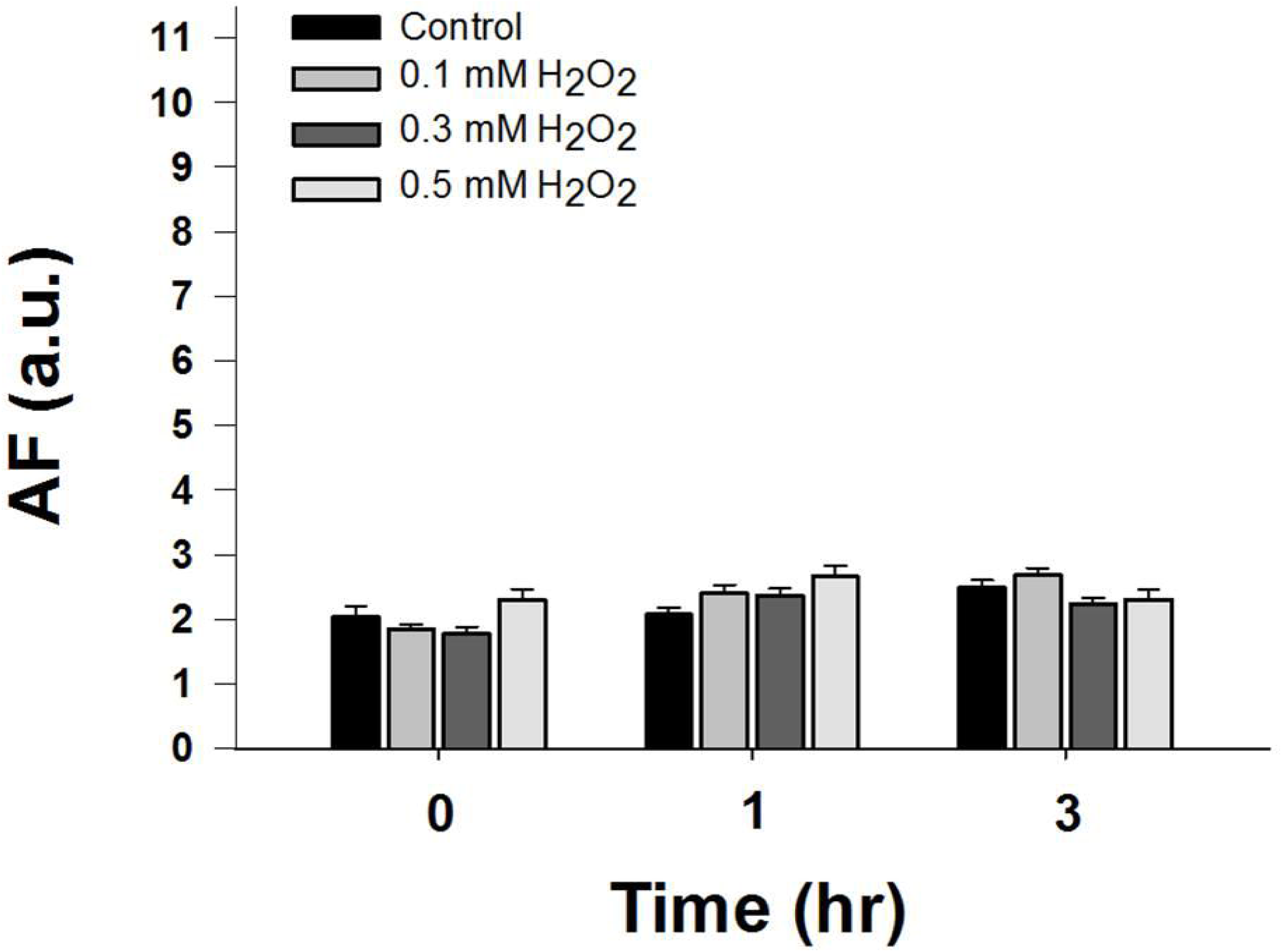
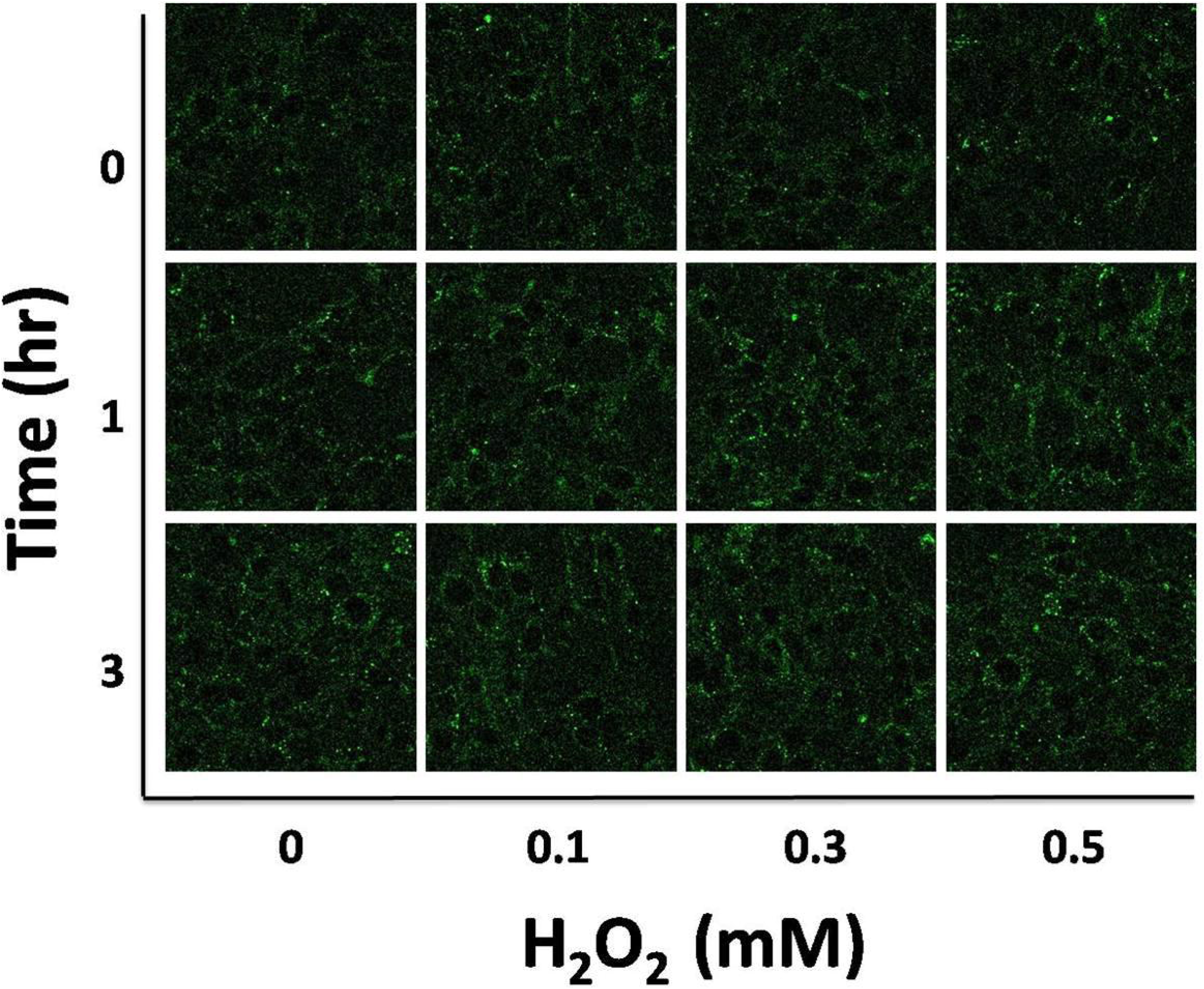
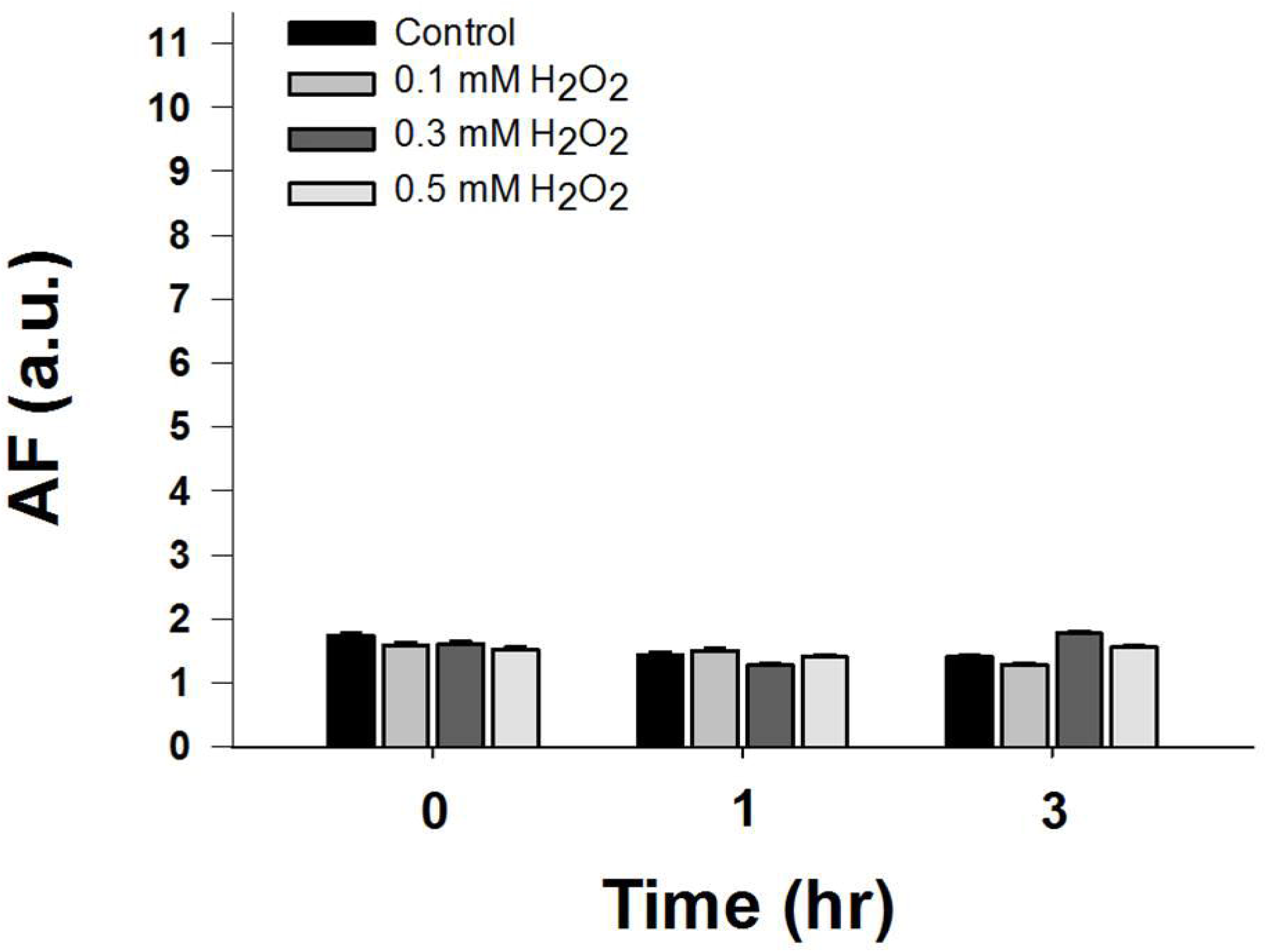
H_2_O_2_ did not affect the green AF of BV2 microglia and PC12 cells. (A) Treatment of BV2 microglia with H_2_O_2_ did not change the AF of the cells. BV2 microglia were treated by 0.1, 0.3, or 0.5 mM H_2_O_2_ for 15 mins. The green AF of the cells was determined under a confocal microscope at 1 hr or 3 hrs after the H_2_O_2_ exposures. The exciting wavelength was 488 nm and the emission wavelength was at the range between 500 and 530 nm. (B) Quantifications of the AF show that H_2_O_2_ H_2_O_2_ did not affect the green AF of BV2 microglia. N = 6. Data were collected from two independent experiments. (C) Treatment of PC12 cells with H_2_O_2_ did not change the AF of the cells. PC12 cells were treated by 0.1, 0.3, or 0.5 mM H_2_O_2_ for 15 mins. The green AF of the cells was determined under a confocal microscope at 1 hr or 3 hrs after the H_2_O_2_ exposures. The exciting wavelength was 488 nm and the emission wavelength was at the range between 500 and 530 nm. (D) Quantifications of the AF show that H_2_O_2_ did not affect the green AF of PC12 cells. N = 6. Data were collected from two independent experiments.

**Supplemental Figure 2.**
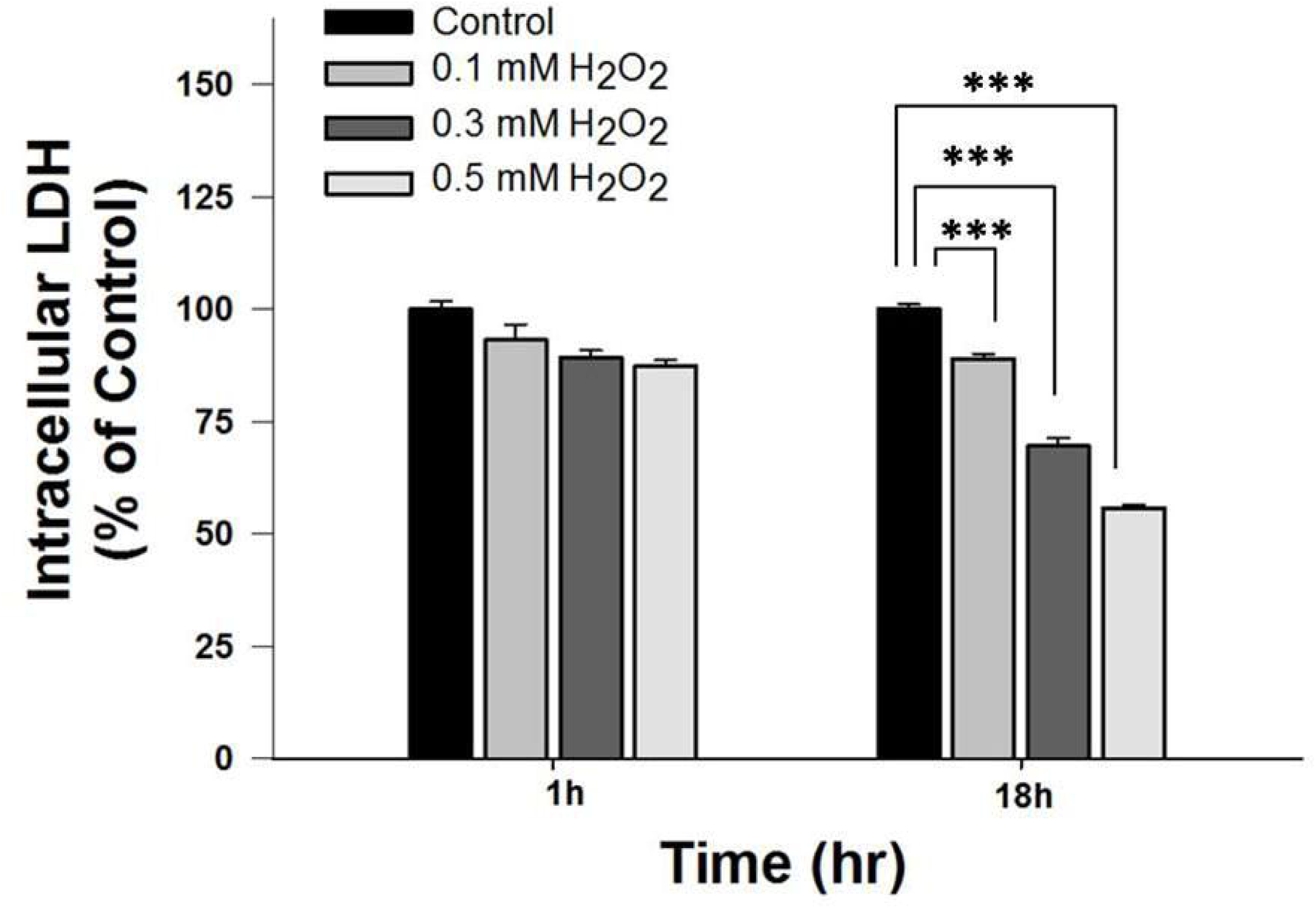
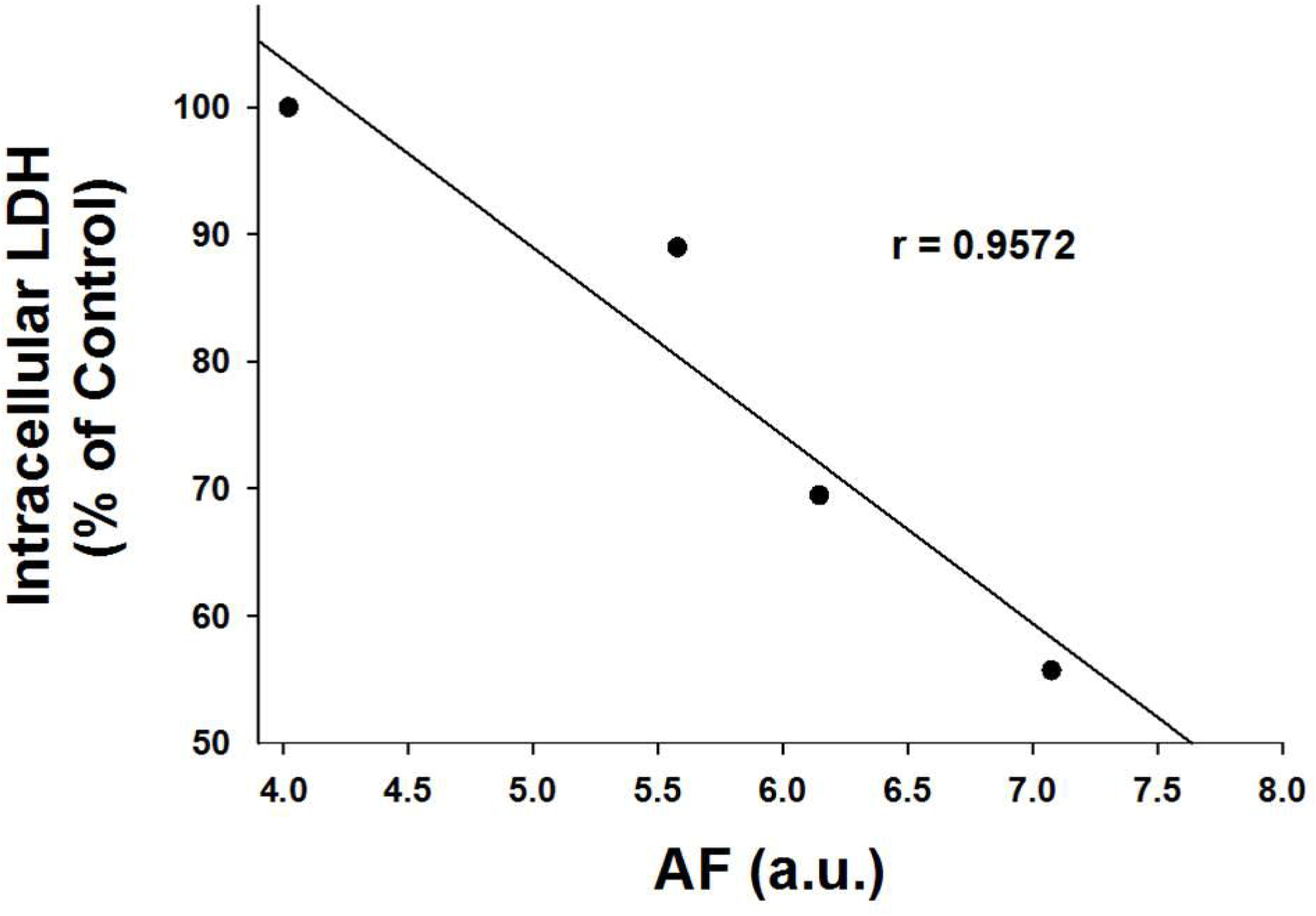
H_2_O_2_-induced AF at 1 hr after H_2_O_2_ exposures is highly negatively correlated with the cell survival of H_2_O_2_-treated HaCaT cells at 18 hrs after H_2_O_2_ exposures. (A) Intracellular LDH assay showed that treatment of HaCaT cells with H_2_O_2_ led to obvious decreases in cell survival at 18 hours after H_2_O_2_ treatment. N = 9. Data were collected from three independent experiments. \**p* < 0.05; \*\*\**p* < 0.001. (B) H_2_O_2_-induced AF at 1 hr after H_2_O_2_ exposures is highly negatively correlated with the cell survival of H_2_O_2_-treated HaCaT cells at 18 hrs after H_2_O_2_ exposures.

## References

1. Afaq F, Syed DN, Malik A, Hadi N, Sarfaraz S, Kweon MH, Khan N, Zaid MA, Mukhtar H. Delphinidin, an anthocyanidin in pigmented fruits and vegetables, protects human HaCaT keratinocytes and mouse skin against UVB-mediated oxidative stress and apoptosis. J Invest Dermatol 127: 222–32, 2007.

2. Byrne AM, Lemasters JJ, Nieminen AL. Contribution of increased mitochondrial free Ca^2+^ to the mitochondrial permeability transition induced by tert-butylhydroperoxide in rat hepatocytes. Hepatology 29: 1523–31, 1999.

3. Cao W, Hong Y, Chen H, Wu F, Wei X, Ying W. SIRT2 mediates NADH-induced increases in Nrf2, GCL, and glutathione by modulating Akt phosphorylation in PC12 cells. FEBS Lett 590: 2241–55, 2016.

4. Cheignon C, Tomas M, Bonnefont-Rousselot D, Faller P, Hureau C, Collin F. Oxidative stress and the amyloid beta peptide in Alzheimer’s disease. Redox Biol 14: 450–464, 2018.

5. Chen H, Wang Y, Zhang J, Ma Y, Wang C, Zhou Y, Gu H, Ying W. NAD^+^-Carrying Mesoporous Silica Nanoparticles Can Prevent Oxidative Stress-Induced Energy Failures of Both Rodent Astrocytes and PC12 Cells. PLoS One 8: e74100, 2013.

6. Huang KF, Ma KH, Liu PS, Chen BW, Chueh SH. Ultraviolet B irradiation increases keratin 1 and keratin 10 expressions in HaCaT keratinocytes via TRPV1 activation and ERK phosphorylation. Exp Dermatol 26: 832–835, 2017.

7. Hurrle S, Hsu WH. The etiology of oxidative stress in insulin resistance. Biomed J 40: 257–262, 2017.

8. Li P, Stetler RA, Leak RK, Shi Y, Li Y, Yu W, Bennett MVL, Chen J. Oxidative stress and DNA damage after cerebral ischemia: Potential therapeutic targets to repair the genome and improve stroke recovery. Neuropharmacology, 2017.

9. M Zhang DTM, H He, Y Li, W Yan, W Yan, Y Zhu, W Ying. UV-Induced Keratin 1 Proteolysis Mediates UV-Induced Skin Damage. bioRxiv, 2017.

10. Ma Y, Cao W, Wang L, Jiang J, Nie H, Wang B, Wei X, Ying W. Basal CD38/cyclic ADP-ribose-dependent signaling mediates ATP release and survival of microglia by modulating connexin 43 hemichannels. Glia 62: 943–55, 2014.

11. Ma Y, Jiang J, Wang L, Nie H, Xia W, Liu J, Ying W. CD38 is a key enzyme for the survival of mouse microglial BV2 cells. Biochem Biophys Res Commun 418: 714–9, 2012.

12. Moran C, Munch G, Forbes JM, Beare R, Blizzard L, Venn AJ, Phan TG, Chen J, Srikanth V. Type 2 diabetes, skin autofluorescence, and brain atrophy. Diabetes 64: 279–83, 2015.

13. Moravcova M, Libra A, Dvorakova J, Viskova A, Muthny T, Velebny V, Kubala L. Modulation of keratin 1, 10 and involucrin expression as part of the complex response of the human keratinocyte cell line HaCaT to ultraviolet radiation. Interdiscip Toxicol 6: 203–8, 2013.

14. Scanlon JM, Aizenman E, Reynolds IJ. Effects of pyrroloquinoline quinone on glutamate-induced production of reactive oxygen species in neurons. Eur J Pharmacol 326: 67–74, 1997.

15. Sinning C, Westermann D, Clemmensen P. Oxidative stress in ischemia and reperfusion: current concepts, novel ideas and future perspectives. Biomark Med 11: 11031–1040, 2017.

16. Takeuchi Y, Hanaoka N, Hanafusa M, Ishihara R, Higashino K, Iishi H, Uedo N. Autofluorescence imaging of early colorectal cancer. J Biophotonics 4: 490–7, 2011.

17. von Leden RE, Yauger YJ, Khayrullina G, Byrnes KR. Central Nervous System Injury and Nicotinamide Adenine Dinucleotide Phosphate Oxidase: Oxidative Stress and Therapeutic Targets. J Neurotrauma 34: 755–764, 2017.

18. Wang H, Joseph JA. Quantifying cellular oxidative stress by dichlorofluorescein assay using microplate reader. Free Radic Biol Med 27: 612–6, 1999.

19. Wu D, Liu B, Yin J, Xu T, Zhao S, Xu Q, Chen X, Wang H. Detection of 8-hydroxydeoxyguanosine (8-OHdG) as a biomarker of oxidative damage in peripheral leukocyte DNA by UHPLC-MS/MS. J Chromatogr B Analyt Technol Biomed Life Sci 1064: 1–6, 2017.

20. Yang C, Ling H, Zhang M, Yang Z, Wang X, Zeng F, Wang C, Feng J. Oxidative stress mediates chemical hypoxia-induced injury and inflammation by activating NF-kappab-COX-2 pathway in HaCaT cells. Mol Cells 31: 531–8, 2011.

